# Screening of natural *Wolbachia* infection in mosquitoes (Diptera: Culicidae) from Cape Verde Islands

**DOI:** 10.1101/2022.11.25.517960

**Authors:** Aires Januário Fernandes da Moura, Vera Valadas, Silvania Da Veiga Leal, Carla A. Sousa, João Pinto

## Abstract

**Background:** *Wolbachia pipientis* is an endosymbiont bacteria that induce cytoplasmic incompatibility and inhibit arboviral replication in mosquitoes. This study aimed at estimating the prevalence and genetic diversity of *Wolbachia* in different mosquito (Diptera: Culicidae) species from Cape Verde.

**Methods:** Mosquitoes were collected in six islands of Cape Verde using dippers/pipettes, BG-sentinel® traps, CDC light traps, and dorsal aspirators. Samples were identified to species using morphological keys and PCR-based molecular assays. *Wolbachia* was detected by amplifying a fragment of the surface protein gene (*wsp*). Multilocus sequence typing (MLST) was performed with five housekeeping genes (*coxA, gatB, ftsZ, hcpA and fbpA*) and the *wsp* hypervariable region (*HVR*) for strain identification. Identification of *w*Pip groups (*w*Pip-I to *w*Pip-V) was performed using PCR-RFLP assay on the ankyrin-domain gene *pk1*.

**Results:** Nine mosquito species were collected, including the major vectors *Aedes aegypti, Anopheles arabiensis, Culex pipiens s*.*s*. and *Culex quinquefasciatus. Wolbachia* was detected in *Cx. pipiens s*.*s*. (100% prevalence), *Cx. quinquefasciatus (98*.*3%), Cx. pipiens/quinquefasciatus* hybrids (100%) and *Culex tigripes (100%)*. Results from MLST and *wsp* hypervariable region typing showed that *Wolbachia* from *Cx. pipiens s*.*l*. belong to Sequence Type 9, *w*Pip clade and supergroup B *Wolbachia*. Phylogenetic analyses indicate that *Wolbachia* isolated from *Cx. tigripes* belongs to Supergroup B but integrates a distinct clade from *w*Pip with no attributed MLST profile. PCR-RFLP revealed *w*Pip-II, *w*Pip-III and *w*Pip-IV groups in *Culex pipiens s*.*l. w*Pip-IV was the dominant group, while *w*Pip-II and *w*Pip-III were restricted to Maio and Fogo islands, respectively.

**Conclusion:** Our study showed a high prevalence and diversity of *Wolbachia* in *Cx pipiens s*.*l*. from Cape Verde islands and, to the best of our knowledge, is the first to detect *Wolbachia* in *Cx. tigripes*, being represented in this species by a previously undescribed MLST Sequence Type.

## Background

*Wolbachia pipientis* (Alphaproteobacteria, Rickettsiales) is an obligate intracellular gram-negative bacteria and proteobacterial symbiont found in a variety of invertebrates, including insects, crustaceans, arachnids, and filarial nematodes (1). Currently, the *Wolbachia* genus is subdivided into 17 supergroups (A-F; H-Q and S) and most species known belong to supergroups A and B (2).

*Wolbachia* is transmitted vertically through host eggs and can influence longevity and reproduction, including feminisation, parthenogenesis and incompatibility between the female and male sex cells (3). The best known phenotype induced by *Wolbachia* in arthropods is cytoplasmic incompatibility (CI). It occurs when males harbouring *Wolbachia* are crossed with uninfected females or between individuals infected with incompatible strains (4,5). The generally accepted model stipulates that cytoplasmatic incompatibility results from a *Wolbachia* “modification” factor (mod; toxin) in the sperm that blocks early embryogenesis, and a *Wolbachia* “rescue” factor (resc; antitoxin) produced in the oocyte that allows the diploid zygote to develop if the cross is compatible (6,7).

Besides cytoplasmic incompatibility, *Wolbachia* can inhibit viral replication in mosquitoes, including Zika, Dengue, West Nile, and Chikungunya arboviruses in *Aedes aegypti* (8,9). Other studies also suggest inhibition of pathogens such as *Plasmodium falciparum* in *An. stephensi* and *An. gambiae* and West Nile Virus in *Cx. quinquefasciatus* (1,10,11). These abilities make *Wolbachia* a promising tool against mosquito-borne diseases and possibly an alternative to conventional vector control programmes using insecticides. In fact, the release of males harbouring incompatible *Wolbachia* into target populations has successfully decreased reproduction by sterilisation (12,13). The release of *Ae. aegypti* transfected with the *Wolbachia w*Mel strain (derived from *Drosophila melanogaster*) led to the establishment of *Ae. Aegypti* populations infected with *Wolbachia* and a proven decrease in dengue incidence in Australia (14) and Malaysia (15).

Cape Verde is threatened by several species of vector mosquitoes, including *Ae. aegypti, An. arabiensis, Cx. quinquefasciatus, and Cx. pipiens s*.*s*. (16). Integrated vector control strategies are mainly directed against *An. arabiensis* and *Ae. aegypti*, using chemical insecticides, diesel, and biological control with *Gambusia* sp. fish (17). However, despite control efforts, the country had its first dengue epidemic in 2009, followed by an outbreak of Zika in 2015-2016 (18) and a malaria outbreak in 2017 (19).

There is no data on the genetic diversity of *Wolbachia* infecting mosquitoes (Diptera: Culicidae) from Cape Verde Islands. This knowledge would be a first step for the design and implementation of programs to suppress mosquito populations through cytoplasmic incompatibility. In this context, the present study aims to detect and genetically characterise *Wolbachia* in populations of Culicidae from Cape Verde.

## Methods

### Study area and sample collection

An entomological survey was carried out in Cape Verde between February and June 2021. Larval and adult mosquito samples were collected on six islands (Santiago, Brava, Fogo, Maio, Santo Antão, and Boavista (Figure 1), using BG-sentinel and CDC light traps, dorsal aspirators, dippers and pipettes. All collection sites were geo-referenced with a portable Global Positioning System (GPS) device (Garmin eTrex® 10).

**Figure 1.**
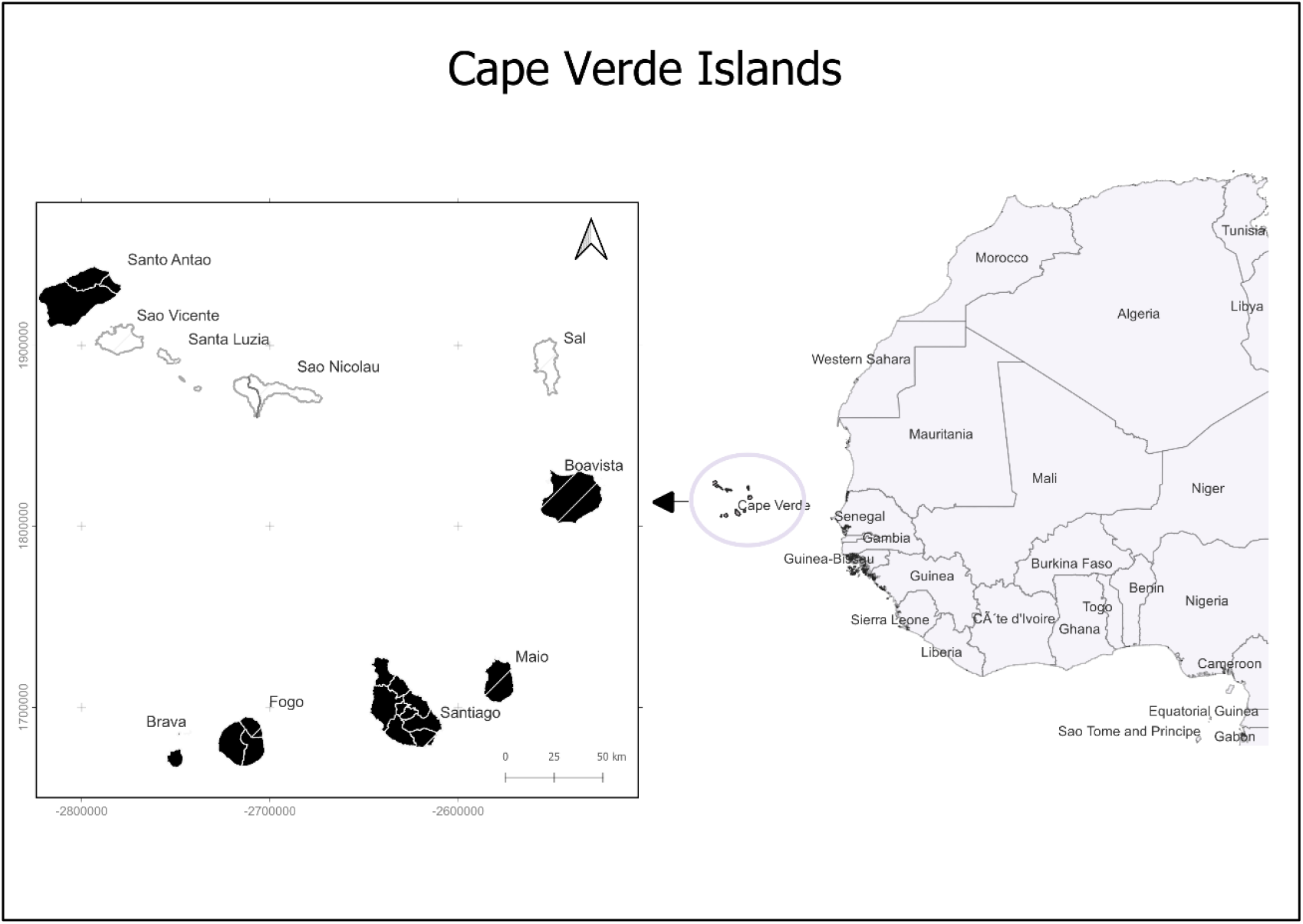
Map of the North Atlantic region showing the geographic location of Cape Verde Islands. Mosquito samples were collected on the islands of Santo Antão, Boavista, Maio, Santiago, Fogo and Brava (highlighted in black)

Mosquitoes were identified to species/complex using the Ribeiro *et al*. identification key (20) and stored individually in microtubes containing silica gel (for adults) or 80% ethanol (for larvae). For genetic analysis, DNA was extracted from single specimens using CTAB 2% and proteinase K, according to Weeks *et al*. (21).

Species of the *An. gambiae* complex were identified by polymerase chain reaction (PCR) according to Scott *et al*. (22), using primer sequences described in Table S1 (supplemental material). PCR was performed using 12.5 μl of Xpert Taq Plus Mastermix (GriSP), 0.1μM of ME and UN primers, 0,05 μM of GA primer, and 0,15 μM of AR primer, plus 1μl of template DNA and water to a final volume of 25 μl.Cycling conditions were as follows: 1 cycle at 95ºC for 5min, 30 cycles at 94ºC for 30s, 50ºC for 30s and 72ºC for 30s; and a final cycle of 72°C for 5min.

For *Culex pipiens* complex, specimens were identified to species by PCR amplification of acetylcholinesterase-2 (*ace-2*) gene sequences using primers described by Smith & Fonseca (23) (Table S1). PCR was performed using 12.5 μl of Xpert Taq Plus Mastermix (GriSP), 0.4μM of ACEquin and B1246 primer, 0,2 μM of ACEpip primer, 1μl of template DNA and water to a final volume of 25 μl. Cycling conditions were performed as follows:1 cycle at 94ºC for 5min, 35 cycles at 94ºC for 30s, 55ºC for 30s, 72ºC for 1min and one cycle at 72ºC for 5min.

Whenever necessary, morphological identification of species other than the above was supported with sequencing of a 710bp fragment of cytochrome c oxidase subunit 1 mitochondrial gene (COI) with primers LCOI1490_F1 and HCOI2198_R1 (Table S1), according to Folmer *et al*. (24). PCR was performed using 1X PCR buffer, 2mM MgCl2, 0.2mM dNTPs, 1U Taq polymerase (Robust HotStart PCR Kit, KAPABIOSYSTEMS), 0.5μM of each primer, 2μl of DNA template, and water to a final volume of 20 μl. Cycling conditions were: initial denaturation at 94°C for 3min; 40 cycles at 94°C for 50s, annealing at 45°C during 30s and 72°C for 1min; and final elongation at 72°C for 5 min.

### Screening of *Wolbachia*

*Wolbachia* detection in mosquito samples was performed by amplifying a 610 bp region of the *Wolbachia* Surface Protein gene (*wsp*), using primers 81F and 691R (Table S2) described by Zhou *et al*. (25). The amplification reaction comprised 12.5 μl of Xpert Taq Plus Mastermix (GriSP), 0.4μM of each primer, 1μl of template DNA and water to a final volume of 25 μl. Cycling conditions were:1 cycle at 95ºC for 3min, 35 cycles at 95ºC for 1min, 55ºC for 1min, 72ºC for 1min and one cycle at 72ºC for 10min. All PCR products from the assays above described were analysed by electrophoresis on a 1.5% agarose gel stained with Green Safe Premium (NZYTech).

### *Wolbachia* MLST and *wsp* typing

*Wolbachia* genotyping was performed through amplification and sequencing of five MLST loci (*gatB, coxA, hcpA, ftsZ, fbpA)* and the *wsp* hypervariable region (26,27). The primer pairs for each locus and the size of amplified products are shown in Table S3 (supplemental materials).

PCR was performed using 1X PCR buffer, 0.2mM dNTPs, 1.5mM MgCl2, 0.5U Taq polymerase (Robust HotStart PCR Kit, KAPABIOSYSTEMS), 1μM of each primer, 2μl of DNA template, and water to a final volume of 40 μl. Cycling conditions were: initial denaturation at 94°C for 2min; 37 cycles at 94°C for 30s, annealing at 54°C (for *hcpA, gatB, ftsZ, and coxA*), and 59°C (*fbpA* and *wsp*), during 45s and 72°C for 1min30s; and final elongation at 72°C for 10 min.

Five microlitres of PCR product from each locus was used in electrophoresis to confirm amplification. The remaining 35μl was purified using Exo/SAP Go-PCR purification kit (GRISP) and sent for direct DNA sequencing at STAB Vida (Oeiras, Portugal) using forward and reverse primers.

*Wolbachia* MLST and hypervariable *wsp* sequences were edited and aligned using BioEdit (version 7.0.9.0) and MEGA software (version 11.0.11). Consensus and concatenated sequences (*gatB, coxA, fbpA, ftsZ, hcpA, wsp HVR*) were queried in *Wolbachia* MLST database (https://pubmlst.org/bigsdb?db=pubmlst_wolbachia_seqdef) for strain characterisation. Sequences were also subjected to the nucleotide Basic Local Alignment Search Tool (BLAST) to verify the similarity with deposited sequences in GenBank (https://blast.ncbi.nlm.nih.gov/Blast.cgi).

Phylogenetic analysis was conducted using the Gamma distributed Tamura 3-parameter nucleotide substitution model, and a Neighbor-Joining tree was generated employing 1000 bootstraps in MEGA software (version 11.0.11).

### Identification of *w*Pip groups by PCR-RFLP

Identification of *w*Pip groups (*w*Pip-I to *w*Pip-V) was performed using PCR-RFLP (restriction fragment length polymorphism) assay based on the ankyrin (ANK) *Wolbachia* marker *pk1* (28–30). A PCR that amplifies a 1300 bp ankyrin domain gene fragment (*pk1*) was performed with primers pk1_For and pk1_Rev (Table S2) (31). The reaction components included 10μl of Xpert Taq Plus Mastermix (GRiSP), 0.4μM of each primer, 2μl of template DNA and water to a final volume of 20μl. Cycling conditions were: 1 cycle at 94ºC for 5 min; 35 cycles at 94ºC for 30s, 52ºC for 30s and 72ºC for 90s; and a final cycle of 72°C for 5min. PCR product was analysed by electrophoresis on a 2% agarose gel stained with Green Safe Premium (NZYTech).

The *pk1* PCR product was digested with restriction enzymes *Taq*αI and *Pst*I to identify different *w*Pip groups (28). Digestion with *Taq*αI was performed with the following reaction mixture: 2 μl of Buffer C (NZYTech), 10 μl of the PCR product, 18 μl of water and 2 μl *Taq*αI enzyme (NZYTech) at 10U/ μl. The mixture was placed in a thermocycler at 65ºC for 1h:30min. The reaction was stopped by adding 0.02mM of EDTA (pH=8) to each tube, and the digestion product was visualised by electrophoresis on a 2% agarose gel. Each allele (*w*Pip group) was detected according to the size of the resulting fragments: allele “a” or “e” (*w*Pip-I or *w*Pip-V; 991, 251, 107 bp); “b” (*w*Pip-III; 669, 665 bp); “c” (*w*Pip-II; 851, 498 bp); “d” (*w*Pip-IV; 497, 251, 107 bp) (28). If alleles “a” or “e” (*w*Pip-I or *w*Pip-V) were present, the two could be differentiated by digesting the *pk1* PCR product with the *Pst*I restriction enzyme. For this purpose, a reaction mixture was prepared with: 2 μl of Buffer A (NZYTech), 12 μl pk1 PCR product, 1μl *Pst*I enzyme (NZYTech) at 10U/ μl and 5 μl of water. The mixture was incubated at 37ºC for 1h, and the reaction stopped by incubating at 80ºC for 20min. Digested DNA fragments were separated on 2% agarose gel electrophoresis. *w*Pip alleles resulting from *pstI* digestion include: “a” (*w*Pip-I; 903, 303, 141 bp); and “e” (*w*Pip-V; 903, 430 bp) (28,29).

Sequencing of *pk1* PCR products was performed to confirm the RFLP profile. For this purpose, the *pk1* PCR product was purified, as described above for the MLST, and sent for direct sequencing using reverse and forward primers. Sequences were subjected to the nucleotide Basic Local Alignment Search Tool (BLAST) and phylogenetic analysis was performed by the Gamma distributed Tamura 3-parameter nucleotide substitution model, and a Neighbor-Joining tree was generated using 1000 bootstraps in MEGA software (version 11.0.11).

## Results

### Mosquito species identification

A total of 1,648 mosquitoes (303 larvae and 1345 adults) were collected. Species identification by morphological characters revealed the presence of *Aedes aegypti* (*N=*663, 40.2%), *Aedes caspius* (*N=*39, 2.4%), *Anopheles gambiae s*.*l*. (*N=*49, 3.0%), *Anopheles pretoriensis* (*N=*275, 16.7%), *Culex pipiens s*.*l*. (*N=*584, 35.4%), *Culex thallassius* (*N=*7, 0.4%), *Culex tigripes* (*N=*3, 0.2%) and *Culiseta longiareolata* (*N=*28, 1.7%).

Ribosomal DNA PCR for identifying species of the *Anopheles gambiae* complex revealed that all collected specimens from this complex belonged to *An. arabiensis*. For the *Cx. pipiens* complex, specimens were identified by *ace-2* PCR as *Cx. pipiens s*.*s*. (*N=*10, 1.7%), *Cx. quinquefasciatus* (*N=*545, 93.3%), and *Cx. pipiens/Cx. quinquefasciatus* hybrids (*N=*29, 5.0%).

### Screening of *Wolbachia*

The *wsp* fragment was amplified only in *Cx. pipiens s*.*s*. (10/10=100% prevalence), *Cx. quinquefasciatus* (536/545= 98.3%), *Cx. pipiens/Cx. quinquefasciatus* hybrids (29/29=100%) and *Cx. tigripes* (3/3= 100%). The remaining species were negative for *Wolbachia*.

### *Wolbachia* MLST and *wsp* typing

We analysed 80 samples that were positive for *wsp* for *Wolbachia* MSLT and *wsp* typing. Allelic profiles resulting from MLST loci and the *wsp* hypervariable region sequencing revealed that *Wolbachia* from *Cx. pipiens s*.*s*., *Cx. quinquefasciatus* and *Cx. pipiens/Cx. quinquefasciatus* hybrids belong to Sequence Type 9, *w*Pip clade and supergroup B *Wolbachia* (Table 1). The same result was obtained from phylogenetic analysis using concatenated sequences of MSLT loci (*coxA, gatB, ftsZ, fbpA, hcpA*) and *wsp* hypervariable region (Figure 2).

**Table 1.** Allelic profile of MLST genes and wsp hypervariable region for different species of Culicidae collected in Cape Verde Islands.)

| Host species | Island (n) | *gatB* | *coxA* | *hcpA* | *ftsZ* | *fbpA* | *wsp* | *HVR1* | *HVR2* | *HVR3* | *HVR4* | Sequence type |
| --- | --- | --- | --- | --- | --- | --- | --- | --- | --- | --- | --- | --- |
| **Cx.**<br>***quinquefasciatus*** | Santiago (n=15) |  |  |  |  |  |  |  |  |  |  | **ST9** |
|  | Brava (n=15) |  |  |  |  |  |  |  |  |  |  |  |
|  | Boavista (n=12) |  |  |  |  |  |  |  |  |  |  |  |
|  | Maio (n= 7) | 4 | 3 | 3 | 22 | 4 | 10 | 10 | 8 | 10 | 8 |  |
|  | Fogo (n=1) |  |  |  |  |  |  |  |  |  |  |  |
|  | S. Antão (n =10) |  |  |  |  |  |  |  |  |  |  |  |
| **Cx. pipiens s.s.** | Maio (n=2) |  |  |  |  |  |  |  |  |  |  |  |
|  | S. Antão (n= 2) | 4 | 3 | 3 | 22 | 4 | 10 | 10 | 8 | 10 | 8 |  |
| **Hybrids pip/qui** | Maio (n=1) |  |  |  |  |  |  |  |  |  |  |  |
|  | Fogo (n=2) | 4 | 3 | 3 | 22 | 4 | 10 | 10 | 8 | 10 | 8 |  |
|  | S. Antão (n=10) |  |  |  |  |  |  |  |  |  |  |  |
| **Cx. tigripes** | Santiago (n=3) | 9 | 182<sup>b</sup> | 12 | 117 | 203<sup>b</sup> | NA<sup>a</sup> | NA<sup>a</sup> | 232 | 222 | 84 | **NA<sup>a</sup>** |

<sup>a</sup> NA represents allelic profile or sequence type not available in the *Wolbachia* MLST database.
<sup>b</sup> Sequences with partial match in the *Wolbachia* MLST database.

**Figure 2.**
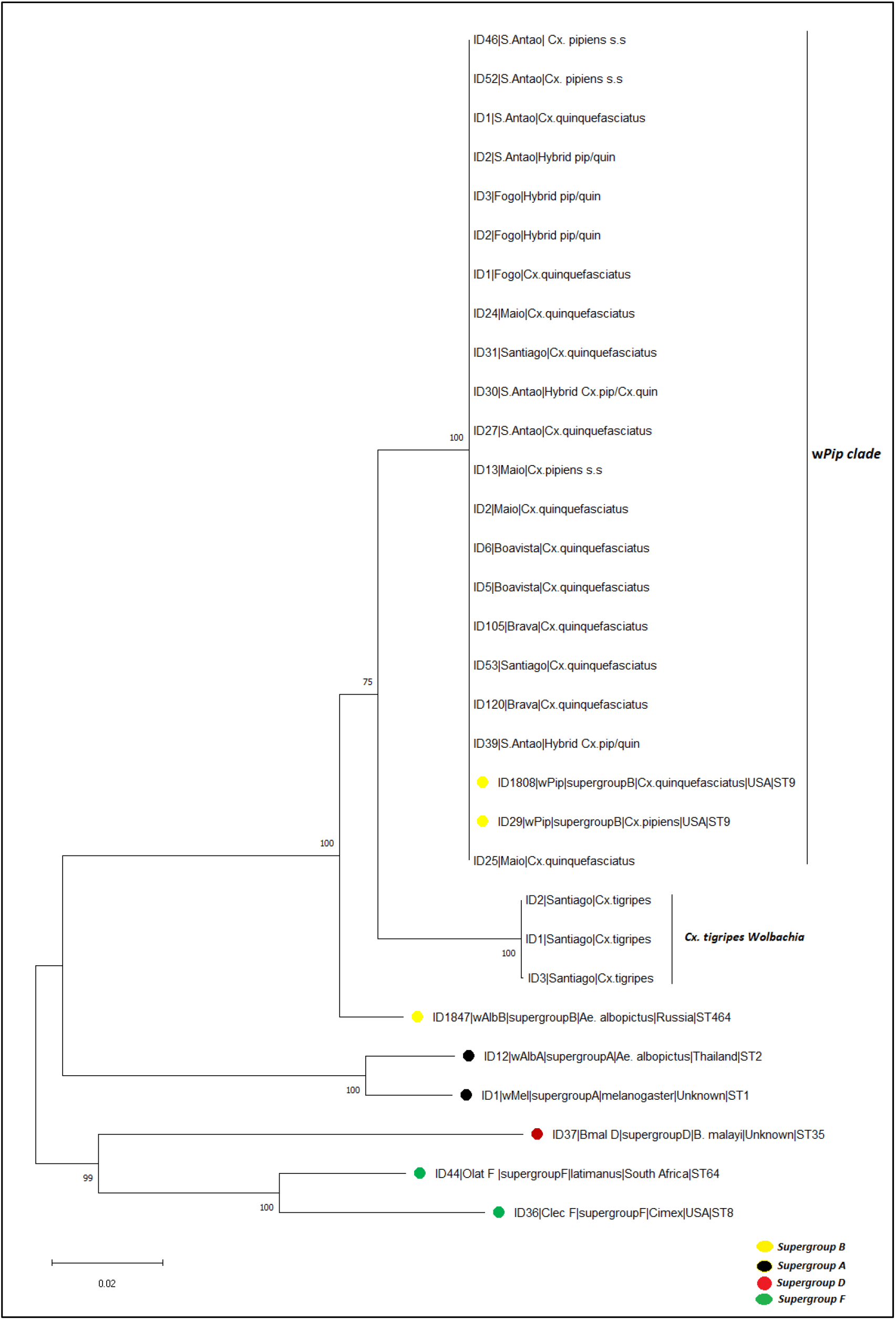
Phylogenetic tree generated from concatenated sequences of MSLT loci (coxA, gatB, ftsZ, fbpA, hcpA) and wsp hypervariable region. Numbers on branches indicate percentage bootstrap support (1000 replicates). Reference sequences were obtained from the Wolbachia MLST database and are marked by full circles. Each Wolbachia supergroup is marked with a different colour. The scale bar indicates the number of substitutions

For *Cx. tigripes*, the allelic profile obtained was unavailable in the MLST database, thus not allowing the determination of a Sequence Type. These MLST sequences were submitted to the MLST database and await attribution of a Sequence Type. The phylogenetic analysis indicates that *Wolbachia* from *Cx. tigripes* belongs to Supergroup B but integrates a distinct clade from *w*Pip (Figure 2).

### *w*Pip groups and their distribution in the archipelago

Results from *pk1* PCR-RFLP showed the occurrence of three different *w*Pip groups in Cape Verde, namely *w*Pip-IV (88.9%), *w*Pip-II (7.4%), and *w*Pip-III (3,7%) (Table 2). The *w*Pip-IV group was detected in *Cx. quinquefasciatus* from five islands (Santiago, Brava, Santo Antão, Maio and Boavista), and also in *Cx. pipiens s*.*s*. from Santo Antão. The *w*Pip-II group was detected only in *Cx pipiens s*.*s*. from Maio, while *w*Pip-III was found exclusively in *Cx. quinquefasciatus* and *Cx. pipiens*/*quinquefasciatus* hybrids from the island of Fogo (Table 2).

**Table 2.** WPip groups detected in Culex pipiens s.l. from Cape Verde Islands according to pk1 PCR/RFLP

| Islands | Species | n | wPip group |
| --- | --- | --- | --- |
| Santiago | *Culex quinquefasciatus* | 15 | wPip- IV |
| Brava | *Culex quinquefasciatus* | 15 | wPip- IV |
| Santo Antão | *Culex pipiens* | 2 | wPip- IV |
|  | *Culex quinquefasciatus* | 10 | wPip- IV |
|  | *Hybrids pipiens/ quinquefasciatus* | 10 | wPip- IV |
| Maio | *Culex pipiens* | 6 | wPip- II |
|  | *Culex quinquefasciatus* | 7 | wPip- IV |
|  | *Hybrids pipiens/ quinquefasciatus* | 1 | wPip- IV |
| Fogo | *Culex quinquefasciatus* | 1 | wPip- III |
|  | *Hybrids pipiens/quinquefasciatus* | 2 | wPip- III |
| Boavista | *Culex quinquefasciatus* | 12 | wPip- IV |

Sequencing of pk1 PCR products confirmed the observed RFLP profiles and similarity with *pk1* sequences deposited in GenBank (Figure 3).

**Figure 3.**
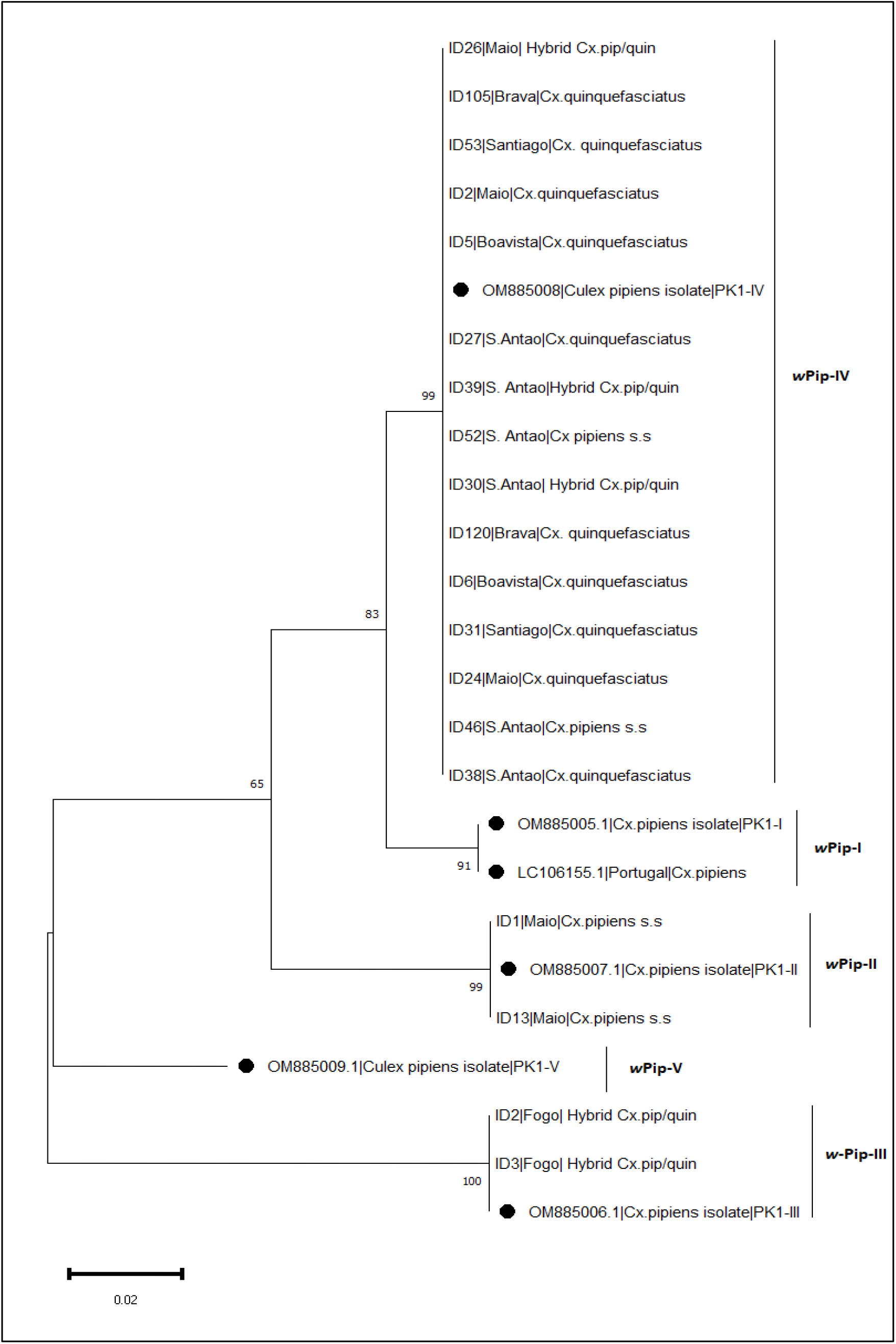
Phylogenetic tree generated from pk1 sequences by Bayesian analysis. Known wPip group pk1 sequences are marked by full circles. Numbers on branches indicate percentage bootstrap support (1000 replicates). The scale bar indicates the number of substitutions.

## Discussion

The MLST and *wsp* typing results revealed that *Wolbachia* from *Cx. pipiens, Cx. quinquefasciatus* and their hybrids belong to the *w*Pip clade and share a monophyletic origin within *Wolbachia* group B. The same results were obtained by Atyame *et al*. (32) and Dumas *et al*. (29) when studying *Wolbachia* genetic diversity from *Cx. pipiens s*.*l*. populations originating from different regions of the world. According to the authors, these findings suggest that *w*Pip strains comprise a recent clade of the *Wolbachia* supergroup B (29,32).

The analysis of the fast-evolving *pk1* gene revealed further diversity within the *w*Pip strain in Cape Verde, indicating the presence of *w*Pip-II, *w*Pip-III and *w*Pip-IV groups in the archipelago. The occurrence of different *w*Pip groups in Cape Verde suggests multiple introduction events in the archipelago. In the past, Cape Verde was a maritime hub between Europe and mainland Africa, and the intense movement of ships may explain the diversity of *w*Pip found in Cape Verde. This result contrasts with that of Southwestern Indian Ocean islands, in which *Wolbachia* infecting *Cx. quinquefasciatus* all belonged to the *w*Pip-I group (13). It is noteworthy that *w*Pip-I was the only group found in mainland Sub-Saharan Africa, South America and Southeast Asia, whereas only *w*Pip-III was detected in North America (29,32). Europe shows the highest diversity, with all five groups of the *w*Pip clade being found in this continent (29). The presence of *w*Pip-II, *w*Pip-III and *w*Pip-IV groups in Cape Verde islands suggests at least three introduction events of *Wolbachia* possibly originating from Europe.However, a North American origin for the *w*Pip-III group in Fogo Island cannot be excluded. Interestingly, the differences found in the genetic composition of the *w*Pip clade among islands agree with the genetic structure of the *Cx. pipiens* complex in Cape Verde. Previous microsatellite-based analysis suggested that *Cx. quinquefasciatus* from Fogo Island may comprise a genetic ancestry cluster distinct from the other islands (33). What was previously considered an admixed *Cx. quinquefasciatus* population in Fogo Island (33) may in fact represent a genetically differentiated population originated from a *w*Pip-III group source population.

The absence of the African *w*Pip-I group from Cape Verde *Cx. quinquefasciatus* is not easily explained. Mainland Africa would be the natural candidate for a source population of *w*Pip-I *Cx. quinquefasciatus* that would have colonized the Cape Verdean islands, as suggested for Southwestern Indian Ocean islands (29). However, *Cx. quinquefasciatus* was predominantly infected by the *w*Pip-IV group. This may suggest that *Cx. quinquefasciatus* from Cape Verde may have derived from a yet to be sampled *w*Pip-IV population of mainland Africa. Another explanation would involve cytoplasmic transfer of *w*Pip-IV from European *Cx. pipiens s*.*s*. to *w*Pip-I *Cx. quinquefasciatus* via hybridization, followed by the replacement of the latter through cytoplasmatic incompatibility (CI). High levels of CI have been reported in crosses between *w*Pip-II and *w*Pip-IV, as well as between *w*Pip-III and *w*Pip-IV infected mosquitoes (7,28). Studies involving experimental crosses would be required to assess CI between *w*Pip-I and *w*Pip-IV and whether this CI would confer an adaptive advantage to *w*Pip-IV infected mosquitoes.

*Wolbachia* was not detected in *Ae. aegypti* from Cape Verde islands, which is consistent with most surveys on this species where no evidence of *Wolbachia* natural infection was found (34–37). Presence of *Wolbachia* in *Ae. aegypti* has been reported only in a few occasions, including those from New Mexico, USA (38) and Kuala Lumpur, Malaysia (39). However, the possibility of *Wolbachia* detection in *Ae. aegypti* being the result of an infection with a *Wolbachia*-carrying nematode or of environmental contamination during field collections, cannot be excluded (35). *Wolbachia* was also not detected in *An. arabiensis* and *An. pretoriensis* from Cape Verde. While this result is in line with most studies that screened for *Wolbachia* in *Anopheles* species (40,41), there have been a few reports on the presence of this endosymbiont in *An. gambiae* and *An. coluzzii* from Mali (42), *An. gambiae* from the Democratic Republic of Congo and *An. coluzzii* in Ghana (43). Shaw *et al*. (44), concluded that *Wolbachia* natural *Anopheles* infections do not induce cytoplasmic incompatibility or sex ratio distortion but show a negative correlation with *Plasmodium* infection, suggesting that *Wolbachia* may interfere with malaria transmission.

This study reports for the first time the presence of *Wolbachia* in *Cx. tigripes*. Phylogenetic analyses indicate that *Wolbachia* isolated from this mosquito belongs to Supergroup B, with no attributed MLST profile. This suggests the presence of a new strain of *Wolbachia* infecting *Cx. tigripes* in Santiago Island.

*Culex tigripes* is the only predatory mosquito in Cape Verde (45), and on the island of Santiago, its larvae are often found in breeding sites associated with *Cx. pipiens s*.*l* species (46). Our results exclude environmental contamination by *Cx. pipiens s*.*l. Wolbachia* since we detected *Wolbachia* in both larvae and an adult male of *Cx. tigripes* (data not shown). More importantly, the concatenated sequences of the MLST *loci* and the *wsp* HVR region clearly showed that the strain detected in *Cx. tigripes* form a monophyletic group separated from the *w*Pip clade. Our phylogenetic analyses also exclude contamination with *Wolbachia* from Supergroup D and F, which are generally found in non-dipterous insects and nematodes (2,47).

## Conclusion

Our study revealed that *Wolbachia* is widespread in *Cx. pipiens s*.*l*. from Cape Verde Islands but absent from other mosquito species, except for *Cx. tigripes*, where a novel *Wolbachia* strain was unveiled. The three distinct *w*Pip groups circulating in *Cx. Pipiens s*.*l*. suggest multiple introduction events in the archipelago, possibly of non-African origin. The finding of a novel *Wolbachia* strain in *Cx. tigripes* may provide an additional candidate to be used in biocontrol approaches. Further studies would be required to isolate this new *Wolbachia* strain to be used in transfection studies with major mosquito vectors in order to assess its potential impact on mosquito reproductive success and vector competence.

## Competing interests

The authors declare that they have no competing interests

## Authors’ contributions

AJM, CAS and JP designed the study; AJM, SLV performed field work;

AJM, VL, performed the molecular laboratory work;

AJM, CAS, SLV, and JP drafted the manuscript.

All authors read and approved the final manuscript.

## Funding information

This work was financed by national funds through FCT – Fundação para a Ciência e Tecnologia, I.P., within the framework of the project ARBOMONITOR (PTDC/BIA-OUT/29477/2017. Aires da Moura was funded by PhD fellowship program of Camões I.P.

## Acknowledgements

We are grateful to the National Institute of Public Health for the laboratory support in Cape Verde, and to technicians from the Ministry of Health, for their assistance in fieldwork.

## Notes

### Competing Interest Statement

The authors have declared no competing interest.

