## Supplementary materials for "Screening of natural *Wolbachia* infection in mosquitoes (Diptera: Culicidae) from Cape Verde Islands"

Table S1- Primer sequences used for molecular identification of mosquitoes species collected in Cape Verde islands.

| Species identification | Primers | Sequences (5’- 3’) | References |
|---|---|---|---|
| ***An. gambiae* complex** | AR (*An. arabiensis*) | AAGTGTCCTTCTCCATCCTA | (Scott et al., 1993) |
| | ME (*An. melas*) | TGACCAACCCACTCCCTTGA | |
| | GA (*An. gambiae*) | CTGGTTTGGTCGGCACGTTT | |
| | UN (*Universal*) | GTGTGCCCCTTCCTCGATGT | |
| ***Culex pipiens* complex** | ACEquin | CCTTCTTGAATGGCTGTGGCA | (Smith & Fonseca, 2004) |
| | ACEpip | GGAAACAACGACGTATGTACT | |
| | B1246s | TGGAGCCTCCTCTTCACGG | |
| **Other species (COI)** | LCOI1490_F1 | GGTCAACAAATCATAAAGATATTG | (Folmer *et al*., 1994) |
| | HCOI2198_R1 | TAAACTTCAGGGTGACCAAAAAATCA | |

Table S2- Primers used for PCR detection of *Wolbachia* and *genotyping w*Pip I-V groups by PCR-RFLP.

| Target | Primers sequences (5'-3') | Size (bp) | References |
|---|---|---|---|
| *wsp* | 81F - TGGTCCAATAAGTGATGAAGAAA | 610 | (Zhou *et al.*1998) |
| | 691R - AAAAATTAAACGCTACTCCA | | |
| *Pk1* | pk1 For - CCACTACATTGCGCTATAGA | 1300 | (Duron *et al.* 2007) |
| | pk1 Rev - ACAGTAGAACTACACTCCTCCA | | |

Table S3- Primers used for *Wolbachia* MLST loci and *wsp* hypervariable region .amplification and sequence analysis.

| Target | Primers sequences (5'-3') | Size (bp) | References |
|---|---|---|---|
| *wsp HVR* | wsp_F1: GTCCAATARSTGATGARGAAAC | 603 | (Baldo *et al*., 2006) (Jolley *et al*., 2018) |
| | wsp_R1: CYGCACCAAYAGYRCTRTAAA | | |
| *gatB* | gatB_F1: GAKTTAAAYCGYGCAGGBGTT | 471 | |
| | gatB_R1: TGGYAAYTCRGGYAAAGATGA | | |
| *coxA* | coxA_F1: TTGGRGCRATYAACTTTATAG | 487 | |
| | coxA_R1: CTAAAGACTTTKACRCCAGT | | |
| *hcpA* | hcpA_F1: GAAATARCAGTTGCTGCAAA | 515 | |
| | hcpA_R1: GAAAGTYRAGCAAGYTCTG | | |
| *ftsZ* | ftsZ_F1: ATYATGGARCATATAAARGATAG | 524 | |
| | ftsZ_R1: TCRAGYAATGGATTRGATAT | | |
| *fbpA* | fbpA_F1: GCTGCTCCRCTTGGYWTGAT | 509 | |
| | fbpA_R1: CCRCCAGARAAAAYYACTATTC | | |
